# Two new qPCR assays for detecting and quantifying the *Aspergillus flavus* and *Aspergillus parasiticus* clades in maize kernels

**DOI:** 10.1101/2024.03.06.583514

**Authors:** A. Leharanger, D. Paumier, B. Orlando, S. Bailly, R. Valade

**Affiliations:** ARVALIS-Institut du Végétal, 91720 Boigneville, France; Mycoscopia, 31470 Fonsorbes, France

**Keywords:** *Aspergillus* section *Flavi*, calmodulin, quick detection, mycotoxins, aflatoxins, food safety

## Abstract

The fungi of *Aspergillus* section *Flavi* mostly produce carcinogenic mycotoxins — aflatoxins (AFs) — of two types: types B and G (AFBs and AFGs). AF are highly hazardous for human and animal health. Their levels in food and feed are therefore highly regulated, with a low acceptable limit for AF content. In France, global warming has led to the detection of AFs in maize harvests since 2015. Mycoflora analyses have identified two species, *A. flavus* (producing AFBs) and *A. parasiticus* (producing both AFBs and AFGs), as responsible for this AF contamination. However, mycoflora analysis is a time-consuming method that cannot readily be applied to large numbers of samples. We propose here an alternative clade-specific functional TaqMan® qPCR method based on the calmodulin gene for distinguishing between the *A. flavus* clade (AfC) and the *A. parasiticus* clade (ApC). We applied this method to 553 maize samples collected in three different harvest years (2018 to 2020). Both clades were detected in about 40% of the samples tested. As expected, we observed significant positive correlations between AFBs and AfC DNA (R² =0.708), and between AFGs and ApC DNA (R² =0.885). This method will be useful for the rapid, simple and cheap characterisation of maize grain contamination with *Aspergillus* section *Flavi*. This method will make it possible to study the relationship between agroclimatic conditions, AF content and species prevalence, to facilitate the anticipation of AF risks due to global warming in France.

## Introduction

The presence of mycotoxins is a matter of great concern due to the significant effects of chronic toxicity due to exposure through food and feed. Aflatoxins (AFs) are the most toxic mycotoxins occurring in host crops infected with *Aspergillus* from section *Flavi* (Flores-Flores et al., 2015). The four primary molecules synthesised by toxigenic fungi from this section are aflatoxins B1, B2 (AFBs), G1, and G2 (AFGs). They can also be metabolised to produce derivatives, such as aflatoxin M1 (AFM1), which may be produced in lactating individuals consuming diets contaminated with aflatoxin B1 (Muaz et al., 2022).

AFs, and more specifically aflatoxin B1 (AFB1) and its derivative AFM1, are considered the most harmful known mycotoxins. AFB1 is a potent natural carcinogen that can cause hepatocarcinoma in humans and is considered by the International Agency for Research on Cancer (IARC) to be a group I carcinogen for both humans and animals (IARC Working Group on the Evaluation of Carcinogenic Risks to Humans, 2012; Wu et al., 2014). This toxin is also immunosuppressive and has been associated with growth impairment in children (Khlangwiset et al., 2011; Meissonnier et al., 2008). AFB1 is generally considered to be a major historical contaminant of crops in tropical and subtropical regions, in which hydrothermal conditions favour both fungal development and toxin production (Diaz, 2005).

AF contamination was not initially considered a significant threat to agricultural production in Europe, due the less favourable climate, and attention was therefore focused on products imported from other countries (EFSA, 2004). Like many countries worldwide, the EU has imposed regulations on the levels of AFs permissible in food, based on the health risk associated with AF consumption (Commission Regulation 2023/915). The maximum permissible content in unprocessed maize destined for human consumption is 2 µg/kg for AFB1 and 4 µg/kg for AFB1, B2, G1 plus G2. Commission Directive 2002/32/EC on undesirable substances in animal feed established maximum contents in feed materials ranging from 5 µg/kg to 50 µg/kg depending on the material and the animal considered.

However, global warming may render climatic conditions more favourable for aflatoxigenic fungi and has already driven the emergence of aflatoxins in maize produced in European areas previously considered to be free of these toxins, such as Romania (Tabuc et al., 2009), Italy (Armorini et al., 2015), Spain (Alborch et al., 2012), Croatia (Pleadin et al., 2015) and France (Bailly et al., 2018). A modelling study has suggested that climate change may greatly increase AF levels in southern Europe (Battilani et al., 2016). Aflatoxins are produced by *Aspergillus* species from *Aspergillus* subgenus *Circumdati* section *Flavi* series *Flavi* (Houbraken et al., 2020). *A. flavus* produces AFBs, whereas *A. parasiticus* can produce both AFBs and AFGs. Based exclusively on the phylogenetic tree for calmodulin in *Aspergillus* (Frisvad et al., 2019; Houbraken et al., 2021; Makhlouf et al., 2019; Susca et al., 2020), the species related to *A. flavus* (*A. aflatoxiformans*, *A. austwickii*, *A. cerealis*, *A. pipericola*, *A. minisclerotigenes* and *A. oryzae*) and those related to *A. parasiticus* (*A. sojae*, *A. novoparasiticus*, *A. arachidicola*, *A. transmontanensis*, *A. sergii*, *A. krugeri* and *A. mottae*) segregate into two distinct groups. For this study, we therefore define the *A. flavus* clade (AfC) as *A. flavus* and all the related species listed above, and the *A. parasiticus*-clade (ApC) as *A. parasiticus* and all the related species listed above. These fungal species are frequent contaminants of many crops, including maize, spices, peanuts, pistachio nuts and cotton seeds, which may be contaminated either before harvest or during storage (Rustom, 1997; Sarma et al., 2017).

In France, Bailly et al., (2018) studied the emergence of aflatoxins and identified *Aspergillus flavus* and *A. parasiticus* as the species responsible for aflatoxin contamination in French maize. They established fungal counts by culture methods and strains from *Aspergillus* section *Flavi* were identified to species level through macroscopic and microscopic examinations. All *Aspergillus* section *Flavi* strains with atypical morphological features were subjected to molecular identification based on the amplification and sequencing of the internal transcribed spacer (ITS), of the beta-tubulin (*BenA*) gene and the calmodulin (*CaM*) gene. This study provided precise information that proved very useful for identifying the *Aspergillus* species responsible for aflatoxin contamination in French maize, but the methods used are time-consuming, costly, and potentially unsuitable for the analysis of large numbers of samples.

Here, we addressed these challenges by developing a rapid, cost-effective quantitative PCR (qPCR) TaqMan® method for specific identification of the *Aspergillus flavus* clade (AfC) and the *A. parasiticus* clade (ApC). We applied these novel methods to the analysis of maize samples collected over a three-year period from French maize harvests and compared the results with those obtained by traditional microbiological and biochemical analyses. We aimed to develop a more efficient and reliable tool for monitoring the emergence of *Aspergillus* species and AF contamination in maize, to facilitate early detection and improve adherence to food and feed safety regulations.

## Materials and Methods

### Sampling strategy

In total, 553 samples were collected from maize in farmers’ fields in France at harvest: 194 samples in 2018, 182 samples in 2019 and 177 samples in 2020 (Fig. 1). The farmers prepared the samples at harvest according to the following instructions: (a) avoid sampling from field margins; (b) avoid the static sampling of grains; (c) sample moving grains during three different periods during the emptying of the combine harvester. In this way, three different subsamples, each weighing at least 1 kg, were collected from the moving grains in the combine harvester during harvest. These three subsamples were then combined to obtain a final sample of at least 3 kg from each farm field.

**Figure 1:**
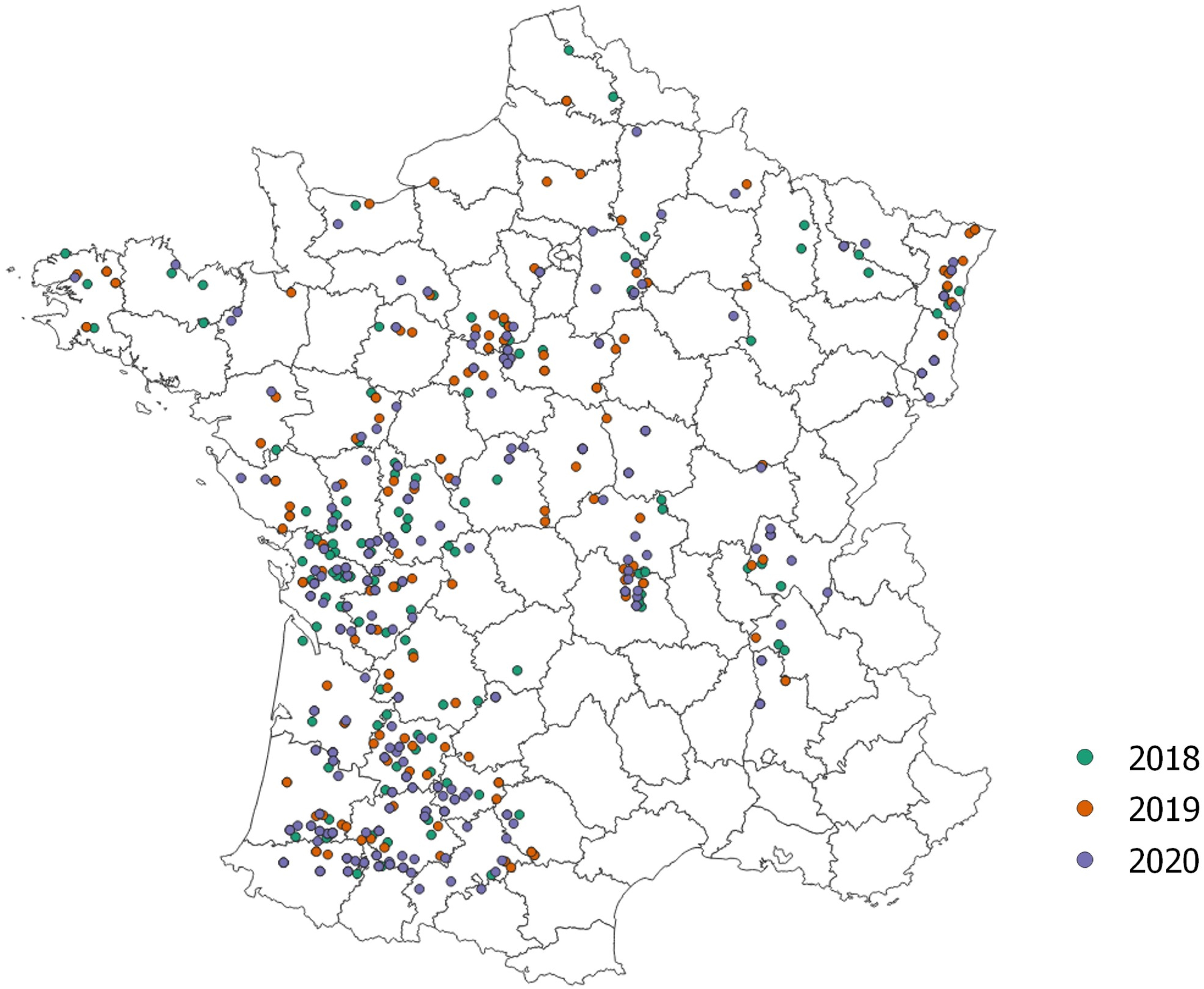
Geographic distribution of the maize farm fields sampled in France by harvest year, for the study period.

All the grain samples were cleaned with a laboratory cleaner and separator (MINI-PETKUS 100 and 200, Rohr, France) to remove all impurities. A 1.5 kg subsample of cleaned and homogeneous grain was then removed, ground in a laboratory hammer mill (FAO, France) fitted with a 1 mm sieve and used for analyses.

### Fungal counts, and identification and quantification of aflatoxins

Fungi were counted and identified as described by Bailly et al. (2018). For this purpose, 20 g of the sample was suspended in 180 mL 0.5% Tween 80. This suspension was subjected to serial 10-fold dilutions, which were plated on both MEA medium and salted MEA (MEA + 6% NaCl). The latter medium was used to identify xerophilic species and to limit the development of pin moulds (Mucorales), which can hinder the correct counting and identification of species with low growth rates. Fungal colonies were counted after three days of culture at 25 ^°^C and confirmed after five days. The limit of detection for fungal counts was 10 CFU/g of sample.

*Aspergillus* section *Flavi* strains were identified to species level by macroscopic and microscopic examinations after five and seven days of culture at 25 ^°^C on MEA and salted MEA. Molecular identification techniques were applied to all *Aspergillus* section *Flavi* strains without the typical morphological features of *A. flavus* and on two *A. flavus* strains as a control. Molecular identification was based on the amplification and sequencing of the internal transcribed spacer (ITS), of the beta-tubulin (*benA*) gene and the calmodulin (*CaM*) gene (Bailly et al., 2018).

Aflatoxins B1, B2, G1 and G2 were analysed by liquid chromatography-tandem mass spectrometry. The limits of detection for AFB1, B2, G1 and G2 were 0.1, 0.1, 0.12 and 0.25 µg/kg respectively, and the corresponding limits of quantification were 0.25, 0.25, 0.25 and 0.5 µg/kg, respectively (Bailly et al., 2018). The results for morphological identification and AF quantification were compared with those of the qPCR method by creating a correlation matrix, to confirm the reliability of the new qPCR system tested.

### Fungal strains

The performance and specificity of the qPCR assays were evaluated with different strains of species from *Aspergillus* section *Flavi* (14 *A. flavus*, 1 *A. aflatoxiformans*, 2 A. *korhogoensis*, 1 *A. minisclerotigenes*, 15 *A. parasiticus*, 1 *A. arachidicola*, 1 *A. sojae*, 1 *A. transmontanensis*, 1 *A. mottae*, 1 *A. sergii*, 1 *A. novoparasiticus*, 1 *A. avenaceus*, 1 *A. bertholletius*, 1 *A. coremiiformis*, 1 *A. leporis*, 1 *A. nomius*, 1 *A. pseudonomius*, and 1 *A. tamarii* isolate) and other pathogenic and non-pathogenic maize fungi (1 *A. niger*, 9 *Fusarium* spp., 2 *Alternaria* spp.). Provenance is reported in Supplementary Table 1 for all the fungal strains used. They were grown in the dark on potato dextrose agar (PDA) at 25 °C with 75% humidity for seven days.

### Genomic DNA extraction

Fungal DNA was extracted from *Aspergillus* sampled from PDA plates by forming pellets of mycelia into 2 mL microtubes and stored at -80 °C for 48 h for freeze-drying. The fungal samples were then ground by two pulses at 30 Hz, for 45 s each, with two sterilised metallic microbeads in an MM400 mixer mill (Retsh). Genomic fungal DNA was extracted with the Macherey Nagel NucleoSpin® Food kit with the corresponding RNAse (20 mg/mL, Macherey Nagel) according to the manufacturer’s protocol. RNAse was added according to the manufacturer’s recommendations. The DNA was eluted in 100 µL elution buffer, in two steps (50 µL each). Fungal DNAs were quantified with a Nanodrop One instrument.

Genomic DNA was extracted from maize samples with the Macherey Nagel NucleoSpin® 96 Food kit (according to the manufacturer’s instructions). A subsample of 100 mg (± 3 mg) of flour was weighed out from each sample. The incubation time was extended to 90 minutes, and RNAse A was added (20 mg/mL, Macherey Nagel). The silica columns were incubated at 37 °C for 20 minutes before the two-step elution process. Nucleic acids were quantified with a Biotek Power Wave XS2 and an adapted spectrometer plate (50 µL).

### Design of specific qPCR primers and TaqMan probes

We ensured that both the assays were of suitable specificity and sensitivity by designing primer pairs and FAM-TAMRA TaqMan probes based on the *CaM* gene encoding calmodulin (listed in Table 1) from 44 *Aspergillus* sequences from the NCBI database (Supplementary Table 2) or published findings (Bailly et al., 2018; Frisvad et al., 2019). Geneious Prime 2023.1.1 was used to align the sequences (Supplementary Figure 1).

**Table 1:**
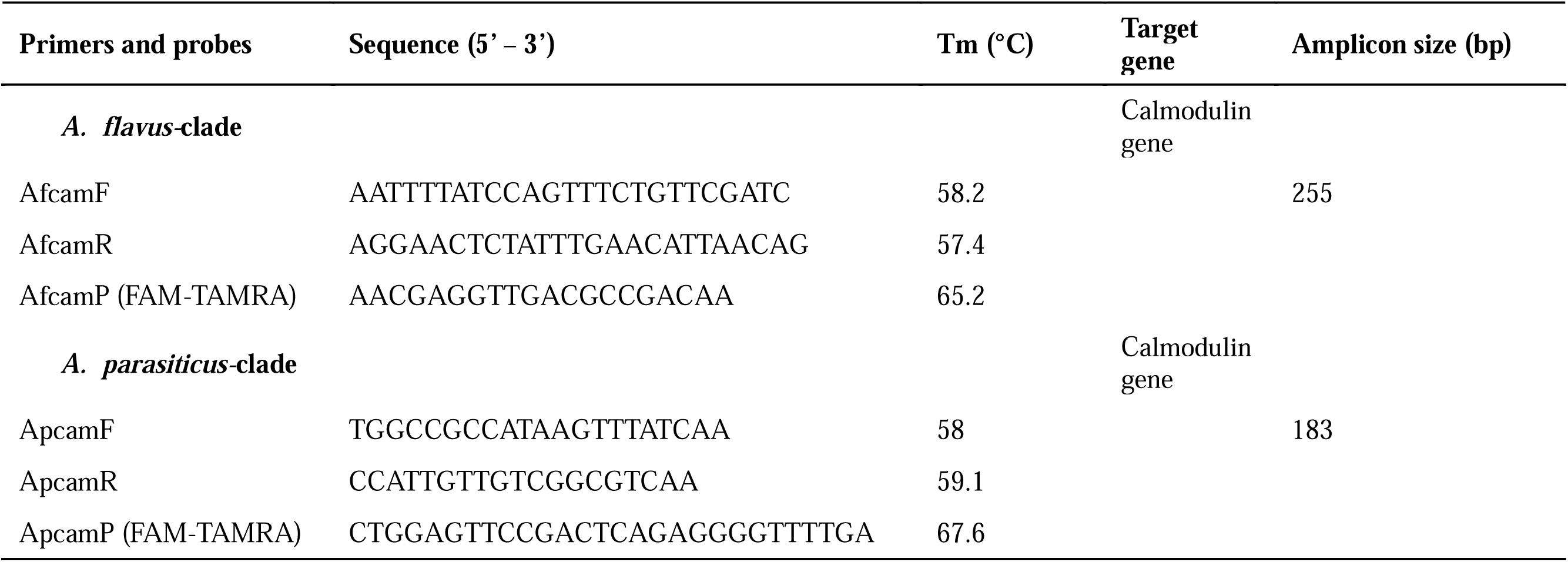
List of sequences of primers and probes designed for *A. flavus*-clade and *A. parasiticus*-clade.

Given the complex nucleotide diversity of the *Aspergillus* species of section *Flavi*, we decided to focus on the design of primers and probes for segregation at clade scale.

Specificity for AfC was achieved by incorporating two different SNPs from ApC in the forward primer AfcamF and four SNPs in the reverse primer AfcamR. Specificity for ApC was achieved by incorporating three SNPs in the forward primer ApcamF and one SNP in the probe ApcamP (Supplementary Figure 1). The specificity of the primers and probes specificity was checked with a BLAST tool online. The formation of secondary structures *in silico* was analysed with OligoAnalyzer 3.1 (https://eu.idtdna.com/calc/analyzer).

### Optimisation of the qPCR assays

Both qPCR assays were optimised to achieve an efficiency of 90% to 105% and an *R²* ≥0.99 for the standard curve. All reactions were performed in Biorad® 96-well plates with the reagent kit for TaqMan^®^ qPCR assays (Eurogentec), in accordance with the manufacturer’s instructions. The optimal primer concentration was 300 nM with MESA Green qPCR^TM^ Mastermix Plus for SYBR^®^ Assays (Eurogentec) and 100 nM for TaqMan^®^ probes. We added 5 µL of sample DNA to obtain a final volume of 25 µL. Each sample was tested in duplicate. Four different concentrations of fungal DNA extract (1 ng.µL^-1^ to 10^-3^ ng.µL^-1^) were analysed in triplicate for the construction of standard curves. The optimal amplification conditions for both primer/probe systems were as follows: 95 °C for 10 min and 40 cycles of 95 °C for 15 s, and 61 °C for 1 min, on a Biorad^®^ CFX Opus 96 instrument. Melting temperature gradients were used to optimise TaqMan® assays.

For each specific clade mix, serial dilutions (1 ng.µL^-1^ to 10^-3^ ng.µL^-1^, five repeats) were used to plot the calibration curve. For the validation of standard curves, Shapiro-Wilk tests were performed, comparing the Cq values with the recalculated Cq values for each of the five dilution replicates (Supplementary Figure 2).

For assays of the specificity of the two primer/probe systems, fungal DNA from each strain (Supplementary Table 1) was diluted to 0.1 ng.µL^-1^ in the final volume. We checked that related *Aspergillus* species did not disturb either of the qPCR assays by spiking *A. flavus* DNA with DNA from 12 fungi from ApC and *A. parasiticus* DNA with DNA from 11 fungi from AfC. Five µL from the ‘pure’ standard diluted DNA was spotted and 4 µL of spiked DNA was added to the mixture. The DNA of each fungal species was present at a concentration of 1 ng.µL^-1^ in the final volume.

Nine serial dilutions, at concentrations ranging from 10 ng.µL^-1^ to 10^-5^ ng.µL^-1^, were used to evaluate the limit of detection (LOD) and the limit of quantification (LOQ), which was calculated by multiplying the LOD by 10. The LOD was reached when 90% of the fungal DNA in the replicate for a standard dilution was detected.

Repeatability was validated for each system by having the same operator generate five standard curves on five different days. The 1 ng.µL^-1^ and 10^-3^ ng.µL (quantification limit) samples of fungal DNA used to generate the standard curve were also deposited on the same plate in duplicate. Their mean Cq value, standard deviation, and the coefficient of variation (CV) were calculated. Reproducibility was checked in a similar manner to repeatability, but with two different operators. Each operator repeated the experiment, five times, on five different days.. Positive and negative controls were included on each qPCR plate.

### qPCR analysis

qPCR data were processed with Biorad® CFX Maestro software. We determined the amount of fungal DNA in the total DNA from the maize sample by using Starting Quantity Sq values (exported from the software) with the following method, with the results expressed as pg.ng_(totalDNA)_^-1^:

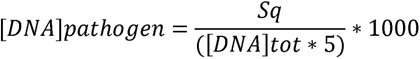

The baseline thresholds were fixed at 30 for AfC and 20 for ApC, to ensure the linearity of the two systems. Maize DNA samples were diluted to 30 ng.µL^-1^ (± 3 ng.µL^-1^).

### Statistical analysis

The correlation matrix was generated and statistical analysis was performed with RStudio software (R version 4.2.2). Input variables were selected according to the data available: section *Flavi* fungal load (CFU.g^-1^), AfC and ApC fungal load (CFU.g^-1^), quantification of aflatoxins B and G (µg.kg^-1^) and quantification of AfC and ApC DNA (pg.ng^-1^). Base 10 logarithmic transformation and scale modification were performed to improve the visualisation of the data distribution on density plots and scatter plots, respectively. Pearson’s correlation coefficients were calculated and their significance assessed.

## Results

### Optimisation and validation of qPCR assays

The primer pairs and probes designed for the two clades had an optimal melting temperature (Tm) of 61 °C (Supplementary Table 3). At this Tm, the standard curves had low Cq values and acceptable efficiency values. Efficiency reached 101.8% (R² = 0.992) for the AfC mix and 98.5% (R² = 0.998) for the ApC mix. The calibration curve showed that efficiency was good, at 98.3% for both the AfC mix and the ApC mix, as was linearity, with values of 0.999 and 1.00, respectively (Supplementary Figure 2). Shapiro-Wilk tests confirmed a good linear relationship between Cq values and recalculated standard values, with *p*-values > 0.05 (Supplementary Figure 2).

In both systems, the LOD was 10^-5^ ng.µL^-1^, giving a LOQ of 10^-4^ ng.µL^-1^. Spiked DNA did not appear to interfere with either of the systems because efficiency (>90%) and reproducibility (>0.990) were good for all curves. Moreover, the amplification curves obtained were similar to that obtained in the absence of spiking.

For the AfC mix, the CVs were 2.64% for the 1 ng.µL^-1^ point and 1.31% for the 10^-3^ ng.µL^-1^ point. For the ApC mix, the CVs were 2.02% and 0.96% for the corresponding points. Both methods were highly repeatable, with CV values of 2.02% and 1.31% for the high- and low-concentration samples, respectively, in the AfC system, and 1.59% and 1.67%, respectively for the ApC system. Both assays were therefore highly reproducible and reliable for the quantification of *Aspergillus* DNA.

### Specificity assays

The main goal of this study was to develop a method capable of distinguishing the species of the *A. flavus* clade from those of the *A. parasiticus* clade. All 58 fungal strains used for the primer/probe specificity tests and the results for their detection with the two qPCR reagent mixes are shown in Table 2. The LOD was 35 (Cq value) for both systems. Based on this threshold, all 14 *A. flavus* strains were successfully detected with the AfC mix. *A. aflatoxiformans*, *A. korhogoensis* and *A. minisclerotigenes* were also detected with the AfC reagent mix. Conversely, no *A. parasiticus* strains or strains of other ApC species were identified using the AfC system. Conversely, no *A. flavus* strains were detected with the ApC-specific primers and probe, and all 14 *A. parasiticus* strains and strains of other ApC species were successfully identified with this mix. *A. coremiiformis* was detected, at the limit of detection, with the ApC mix. No *Fusarium* or *Alternaria spp.* were detected in quantifiable amounts in either assay. The specificity of each primer/probe pair was, therefore, validated.

**Table 2:** Specificity assay for each *Aspergillus* clade reagents mix. Samples are tested in duplicate. “+” = presence; “-” = absence; “LOD” = sample that reaches the LOD (10^-3^ ng.µL^-1^). Out of 35Ct, samples are considered as absent.

| Samples | *A. flavus*-clade system |  | *A. parasiticus*-clade system |  |
| --- | --- | --- | --- | --- |
|  | Cq mean value | Result | Cq mean value | Result |
| ***Aspergillus* spp** |  |  |  |  |
| ***A. flavus*-clade** |  |  |  |  |
| *A. aflatoxiformans* | 24.29 | + | N/A | - |
| *A. flavus* n=14 (min – max) | 24.35 – 28.14 | + | 37.16 – N/A | - |
| *A. korhogoensis* 01 | 24.40 | + | 35.38 | - |
| *A. korhogoensis* 02 | 27.20 | + | N/A | - |
| *A. minisclerotigenes* | 23.82 | + | 35.66 | - |
| ***A. pararasiticus*-clade** |  |  |  |  |
| *A. arachidicola* | N/A | - | 28.34 | + |
| *A. mottae* | N/A | - | 25.45 | + |
| *A. novoparasiticus* | N/A | - | 24.34 | + |
| *A. parasiticus* n=15 (min – max) | N/A | - | 23.47 – 28.77 | + |
| *A. sojae* | 34.34 | LOD | 26.56 | + |
| *A. transmontanensis* | N/A | - | 31.13 | + |
| *A. sergii* | N/A | - | 24.41 | + |
| **Other clades, section or species** |  |  |  |  |
| *A. avenaceus* | N/A | - | N/A | - |
| *A. bertholletius* | N/A | - | N/A | - |
| *A. coremiiformis* | 34.77 | LOD | 28.24 | + |
| *A. leporis* | N/A | - | N/A | - |
| *A. pseudonomius* | N/A | - | N/A | - |
| *A. niger* | N/A | - | N/A | - |
| *A. nomius* | N/A | - | N/A | - |
| *A. tamarii* | N/A | - | N/A | - |
| ***Fusarium* spp** |  |  |  |  |
| *F. avenaceum* | N/A | - | N/A | - |
| *F. graminearum* | N/A | - | N/A | - |
| *F. poae* | N/A | - | N/A | - |
| *F. proliferatum* | N/A | - | N/A | - |
| *F. sporotrichoides* | N/A | - | N/A | - |
| *F. subglutinans* | N/A | - | N/A | - |
| *F. temperatum* | N/A | - | N/A | - |
| *F. tricinctum* | N/A | - | N/A | - |
| *F. verticillioides* | N/A | - | N/A | - |

**Table 2:** Specificity assay for each *Aspergillus* clade reagents mix. Samples are tested in duplicate. “+” = presence; “-” = absence; “LOD” = sample that reaches the LOD ( $10^{-3}$ ng. μL $^{-1}$ ). Out of 35Ct, samples are considered as absent.
|  |  |  |  |  |
| --- | --- | --- | --- | --- |
| *A. section porri* | N/A | - | N/A | - |
| *A. section alternaria* | N/A | - | N/A | - |

### Analysis of maize samples

Once the two qPCR assays had been optimised and validated, we used them to test maize samples. In total, 553 samples from three harvests (2018, 2019 and 2020) were analysed with the two qPCR systems. For mycoflora and fungal DNA quantification results, a fungal profile was assigned by each method (Table 3). For example, if positive PCR results were obtained with the AfC system but not with the ApC system, the sample was classified as AfC. Conversely, if the sample yielded positive results only with the ApC system, it was classified as ApC. Samples yielding positive results with both systems were annotated as ’copresence.’ This classification approach made it possible to obtain a comprehensive characterisation of the fungal profile of each sample based on qPCR and mycoflora analyses.

**Table 3:** Proportion of absence, presence and copresence of fungal profile using mycoflora or qPCR results (a); Proportion of absence, presence and copresence of fungal profile based on qPCR profile and aflatoxins profile over the three years (b).

| Year | Detection method | Absence (%) | AfC profile (%) | ApC profile (%) | Copresence (%) |
| --- | --- | --- | --- | --- | --- |
| **2018** | *Mycoflora* | 46.4 | 46.9 | 2.6 | 4.1 |
|  | *qPCR* | 73.2 | 23.2 | 0.5 | 3.1 |
|  | *Mean qPCR (pg.ng<sup>-1</sup>)* | *n = 142* | 0.01172 (*n = 45*) | 0.00137 (*n = 1*) | 0.15205 (*n = 6*) |
|  | *Min – Max qPCR (pg.ng<sup>-1</sup>)* | / | 0.00077 – 0.15682 | / | 0.00707 – 0.38869 |
| **2019** | *Mycoflora* | 44.0 | 51.1 | 1.1 | 3.8 |
|  | *qPCR* | 61.0 | 28.0 | 1.6 | 9.3 |
|  | *Mean qPCR (pg.ng<sup>-1</sup>)* | *n = 111* | 0.00927 (*n = 51*) | 0.00348 (*n = 3*) | 0.04858 (*n = 17*) |
|  | *Min – Max qPCR (pg.ng<sup>-1</sup>)* | / | 0.00078 – 0.13552 | 0.00150 – 0.00601 | 0.00433 – 0.27722 |
| **2020** | *Mycoflora* | 21.0 | 65.0 | 0.6 | 13.6 |
|  | *qPCR* | 44.1 | 37.9 | 1.1 | 16.9 |
|  | *Mean qPCR (pg.ng<sup>-1</sup>)* | *n = 78* | 0.04538 (*n = 67*) | 0.00435 (*n = 2*) | 0.19438 (*n = 30*) |
|  | *Min – Max qPCR (pg.ng<sup>-1</sup>)* | / | 0.00071 – 2.25000 | 0.00424 – 0.00446 | 0.00416 – 1.79450 |
| **All three years** | *Mycoflora* | 37.4 | 54.1 | 1.5 | 7.1 |
|  | *qPCR* | 59.9 | 29.5 | 1.1 | 9.6 |
|  | *Mean qPCR (pg.ng<sup>-1</sup>)* | *n = 331* | 0.02492 (*n = 163*) | 0.00342 (*n = 6*) | 0.14282 (*n = 53*) |
|  | *Min – Max qPCR (pg.ng<sup>-1</sup>)* | / | 0.00071 – 2.25000 | 0.00137 – 0.00601 | 0.00416 – 1.79450 |

|  | qPCR |  |  |  |  |
| --- | --- | --- | --- | --- | --- |
|  | Absence (%) | AfC (%) | ApC (%) | Copresence (%) | Total (%) |
| **Aflatoxins** |  |  |  |  |  |
| <LOQ | 59.3 | 27.7 | 1.1 | 4.9 | 92.9 |
| AfB | 0.4 | 1.6 | / | 1.6 | 3.6 |
| AfG | 0.2 | / | / | / | 0.2 |
| AfB & AfG | / | 0.2 | / | 3.1 | 3.3 |

Considering the samples of all three years together, 40.1% (Table 3A) of the 553 samples gave positive results in one or both of the qPCR assays. AfC was detected more frequently (29.5%) than ApC (1.1%). We found that 9.6% of the samples were contaminated with both fungal clades. According to our qPCR results, more than half the samples collected in 2020 were positive (55.9%), whereas the proportion of positive samples was lower in 2018 and 2019 (26.8% and 39% respectively). A similar pattern was observed with the mycoflora data.

The DNA quantification data were compared with the *Aspergillus* section *Flavi* mycoflora quantification data for each year. There was a strong, significant correlation (correlation coefficient = 0.904) between total *Aspergillus* section *Flavi* mycoflora data and the total amount of AfC and ApC DNA in all three years, and this correlation was consistently strong for each individual year considered separately (Figure 2). The correlation between the amount of AfC DNA and the total mycoflora was also very strong and significant (0.988) and there was a significant but moderate correlation between the amount of ApC DNA and the total mycoflora (0.534).

**Figure 2:**
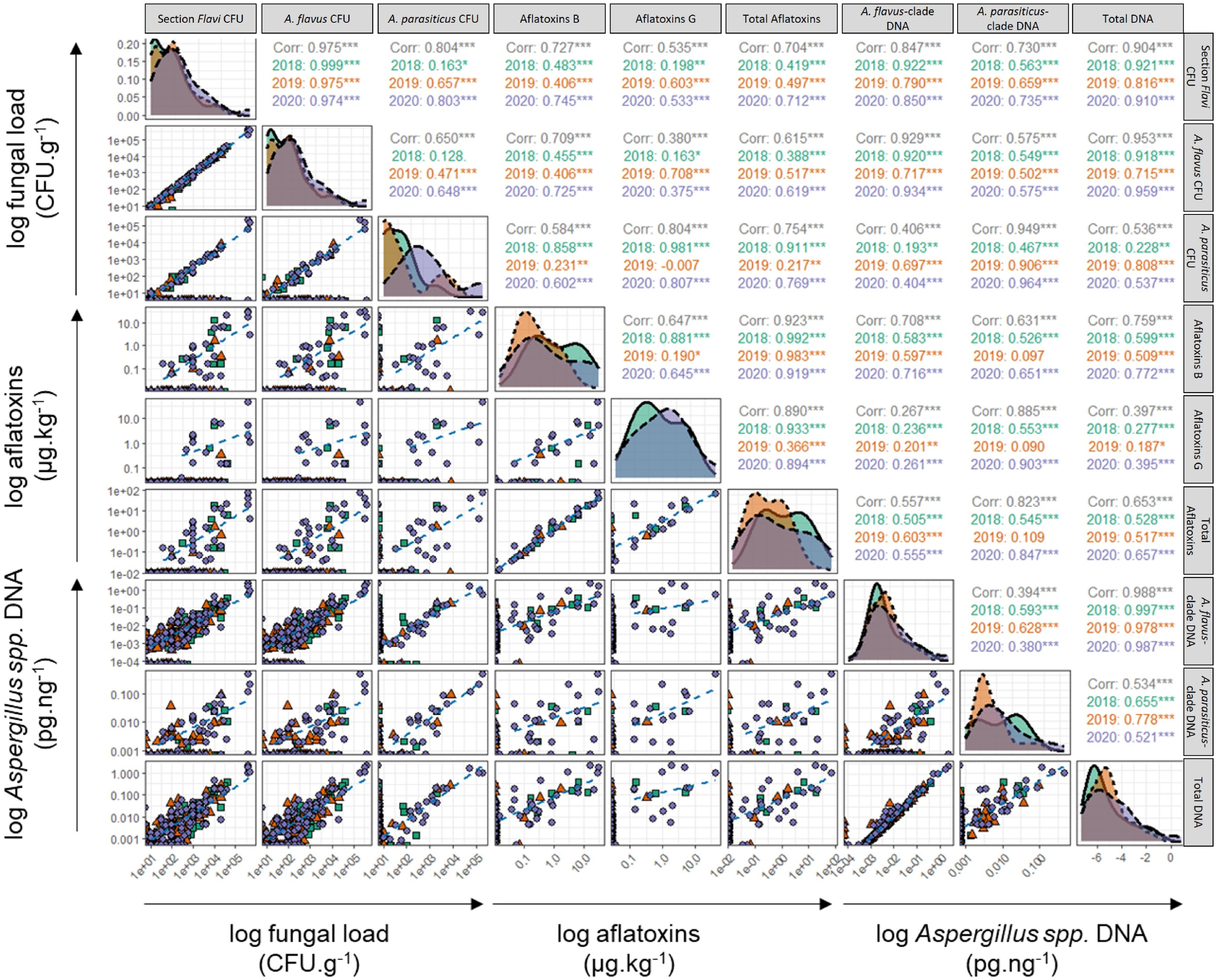
Correlogram for the relationships between fungal load, aflatoxin content and qPCR results. Along the diagonal, density plots: light green area and solid line = 2018, light red area and dotted line = 2019, light blue area and dashed line = 2020. To the right of the diagonal, Pearson correlation coefficients and their significance, the black text indicates the correlation coefficient for all three years considered together. To the left of the diagonal, scatter plots: light green squares = 2018, light red triangles = 2019, light blue circles = 2020; the light blue dashed line is a smooth trend line for 2020 only. Correlations were calculated without data transformation. Density plots were generated from log-transformed data. α = 0.05, * = *p*-value<0.05, ** *= p*-value<0.01, *** = *p*-value<0.001.

We also acquired aflatoxin quantification results for these three years. Only 7.1% of the total samples gave positive results for AF quantification, with most detections occurring in 2020 (Table 3B). Overall, 6.5% of samples tested positive for both aflatoxins and fungal DNA on qPCR, whereas 0.6% of samples contained quantifiable amounts of aflatoxins but did not test positive for the presence of fungal DNA. In the three years studied, 27.7% of the samples were identified as AfC by qPCR without aflatoxin detection during the aflatoxin quantification process.

A comparison of the amounts of DNA for the two clades with the results of aflatoxin quantification gave consistent results. Indeed, the amount of AfC DNA was correlated with the amounts of the B_1_ and B_2_ aflatoxins together, with coefficients of 0.583 for 2018, 0.597 for 2019, 0.716 for 2020 and 0.708 for all three years considered together. The correlation between the amount of AfC DNA and aflatoxin type G content was weaker than the correlation observed for type B aflatoxins (correlation coefficient: 0.267). The correlation between total aflatoxin content and the amount of AfC DNA was strong and significant. As expected, the coefficients of the correlations between ApC DNA content and the amounts of the B and G aflatoxins were high (0.631 and 0.885, respectively).

## Discussion

Climate change may be driving the increasing prevalence of *Aspergillus* section *Flavi* contamination in European crops in the field and during storage (Bailly et al., 2018).

Improvements in the management of these fungi, which produce mycotoxins hazardous to human and animal health, will require the development of rapid, cost-effective methods for their detection and quantification. The methods widely used for detecting *Aspergillus* section *Flavi* include morphological analysis after the development of colonies on a medium favouring fungal growth (Bailly et al., 2018; Pitt et al., 1983; Pitt and Hocking, 2009; Samson et al., 2010; Varga et al., 2011), but these methods are cumbersome and time-consuming. The development of specific molecular markers, as in this study, should accelerate the identification process and improve our understanding of the epidemiology of these fungi.

We developed a rapid diagnostic method based on two pairs of markers, each targeting a single gene, for the specific qPCR-based quantification of *A. flavus* and *A. parasiticus*. Several genes, such as BenA, and ITS sequences are known to give specific amplification products on PCR (Frisvad et al., 2019; Sardiñas et al., 2011). Susca et al. (2019) introduced an SYBR qPCR system using the calmodulin gene to distinguish between toxigenic fungal species linked to maize ear rot, including a number of *Aspergillus* species. However, the primers developed were unable to differentiate between *A. flavus* and *A. parasiticus*. According to Okoth et al. (2018), the calmodulin genes of different *Aspergillus* species have more SNPs in common than the BenA gene or ITS sequences of *Aspergillus* isolates from maize samples. During primer and probe design, a bioinformatic analysis indicated that it would be difficult to discriminate *A. flavus* or *A. parasiticus* from the rest of section *Flavi* (Frisvad et al., 2019). Indeed, both of the qPCR systems based on the calmodulin gene are clade-specific rather than species-specific because the SNPs distinguishing between *A. flavus* and *A. parasiticus* are also present in several of the species closely related to these two taxa. Moreover, the classification of *Aspergillus* section *Flavi* species is not yet stable, with new species continually being added. For example, two *A. parasiticus* were incorrectly identified in one recent study (Chang, 2021) and 107 accepted *Aspergillus* species were added to the list drawn up by Houbraken et al. (2020) between 2014 and 2020.

We therefore decided to develop a method for distinguishing between the two clades, differentiating between AfC and ApC. The AfC mix detected only AfC members (*A. flavus, A. aflatoxiformans, A. minisclerotigenes, A. korhogoensis*) (Table 2) whereas the ApC mix detected only ApC members (*A. parasiticus, A. novoparasiticus, A. arachidicola, A. mottae, A. sergii, A. transmontanensis, A. sojae*). Except for *A. coremiiformis,* these results are consistent with the primer and probe design (Supplementary Figure 1). *A. coremiiformis* does not produce aflatoxins or kojic acid and little new information about this species has been published since 1978 (Bartoli and Maggi, 1978; Frisvad et al., 2019; Varga et al., 2011). The probability of detecting *A. coremiiformis* in French maize crop is therefore low. Thus, our qPCR assays are clade-specific, repeatable, and reproducible.

Another challenge in pathogen DNA quantification from field or storage samples is detecting only the DNA of interest in the total sample DNA. This requires a very sensitive qPCR assay. Thus, with the aim of developing a rapid, cheap and sensitive qPCR-based method for *Aspergillus* section *Flavi* detection and quantification in maize samples, we decided to make use of TaqMan® probe chemistry, which has been shown to be more specific and sensitive than SYBR® (Navarro et al., 2015). The assays developed here detected 0.00071 pg.ng^-1^ to 2.25 pg.ng^-1^ AfC DNA and 0.00137 pg.ng^-1^ to 0.00601 pg.ng^-1^ ApC DNA (Table 3A). These are relatively low concentrations of DNA, demonstrating the high sensitivity of our qPCR assays.

After optimisation of the two qPCR assays, we analysed 553 maize samples from the 2018 to 2020 harvests for which mycoflora and aflatoxin quantification data were available. The correlations between DNA quantification results with the two qPCR systems and the other two sets of data were encouraging for the three years considered together (Figure 2). Indeed, the coefficient of correlation between ApC and total aflatoxin (AFGs+AFBs) levels reached 0.823 (for the three years together), reflecting the capacity of *A. parasiticus* to produce both types of aflatoxin. Conversely, the coefficient of correlation between AfC DNA levels and AFG content was low (0.267) but non-zero, due to the copresence of AfC and ApC (9.6%, Table 3). This result is consistent with the inability of *A. flavus* to produce AFGs. The presence of species producing AFBs and AFGs belonging to AfC, such as *A. korhogoensis* (Carvajal-Campos et al., 2017) and *A. minisclerotigenes* (Frisvad et al., 2019) may affect the correlation and bias the interpretation. Our results show that contamination with AFBs results principally from the presence of *A. flavus*. However, *A. minisclerotigenes* has already been reported twice in Europe (Bailly et al., 2018; Soares et al., 2012). Caution is therefore required in the interpretation of AfC DNA quantification results, which may underestimate the true AFG content.

ANOVA with Tukey HSD tests revealed that the qPCR results for 2018 and 2019 were similar but that the results for each of these years were significantly different from those for 2020, except for ApC DNA quantification, for which the results obtained in 2019 and 2020 were similar. This finding for tests on naturally infected maize seeds in three different harvest years indicates that the results obtained with our qPCR assays are in line with those for mycoflora and aflatoxin analyses, demonstrating the reliability of our approach. The lack of statistical significance for AFG and ApC DNA quantification may result from the small number of samples in which only the *A. parasiticus* clade was detected. Indeed, levels of contamination with *Aspergillus* spp. were high in 2020, in terms of both the proportion of samples in which both *Aspergillus* clades were present and the mean concentration of fungal DNA (mostly the *A. flavus* clade) in this year. A larger proportion of samples tested positive for AfC by mycoflora analysis than with the qPCR detection method in each year. However, the proportions of samples in which only ApC was detected or in which either or both of the two clades were detected were similar for the two methods. Moreover, in 2019 and 2020, ApC seemed to be more present in copresence with AfC, according to the results of both detection methods (for 2019, 3.8% and 9.3%, for mycoflora analysis and fungal DNA quantification, respectively; for 2020, 13.6% and 16.9%, respectively), but ApC was less frequent than AfC overall. Information about agroclimatic conditions during the three years studied might provide an explanation for this situation. Environmental conditions in 2020 were probably favourable for the growth of *Aspergillus,* leading to the high levels of fungal DNA detected and to the high levels of contamination in mycoflora analysis and AF quantification. *Aspergillus* section *Flavi* interspecies interactions have recently been shown to play a role in the infection of maize, by regulating the mycelial growth or sporulating capacity of each of the species in competition for proliferation (Ching’anda et al., 2021). Moreover, certain kinds of aflatoxigenic *Aspergillus* spp. were found to grow better, with higher rates of sporulation, and to more aflatoxins at higher temperatures, indicating that temperature may favour proliferation of certain species of *Aspergillus* section *Flavi* in certain situations (Ching’anda et al., 2021).

Furthermore, the lower frequency of aflatoxin detection in samples than of fungal DNA detection in our qPCR assays may be accounted for by the differences in sensitivity between these methods. The LC-MS-MS method can detect aflatoxins produced in favourable conditions by aflatoxigenic strains only, whereas the qPCR assays developed here quantify the presence of all strains (toxigenic or atoxigenic) targeted by the markers, even if a toxigenic strain cannot produce aflatoxins due to unfavourable conditions. Moreover, had the LC-MS-MS method been more sensitive (lower LODs), it would have detected more aflatoxins. The two qPCR assays presented here are clade-specific, but they make it possible to detect and quantify species from *Aspergillus* section *Flavi* in maize kernels with a high sensitivity.

This approach could be enhanced by the development of a multiplex qPCR system, based on TaqMan® chemistry (Navarro et al., 2015), which would reduce experimental time and facilitate quantification. It would also be of interest to establish a digital PCR protocol to improve sensitivity. Unlike qPCR, dPCR provides absolute quantification, avoiding the need for standard ranges and for the blocking of qPCR inhibitors (Huggett et al., 2015; Sidstedt et al., 2020). The use of dPCR has been increasing in recent years and this technique has already been developed for the detection and identification of the commercialised biocontrol strain of *A. flavus,* AF36, for assessment of its competitiveness with aflatoxigenic *A. flavus* strains (Hua et al., 2018; Schamann et al., 2022).

In conclusion, we have developed and validated two qPCR assays for the rapid identification and quantification of *Aspergillus* from the *parasiticus* and *flavus* clades in maize kernels. This method will also be useful for analyses on many other crops. These methods are complementary to microbiological analyses and methods of aflatoxin detection and quantification. We analysed numerous samples from three different maize harvests and detected the presence of low levels of *Aspergillus* section *Flavi* in French maize harvests. This study provides a first database, which can be used in conjunction with mycoflora, aflatoxin quantification and DNA quantification data from previous and future harvests. Agronomic and climatic data may improve our understanding of the dynamics of contamination leading to the establishment of *Aspergillus* section *Flavi* in France, and of links to mycotoxin production, thereby facilitating the development of disease management strategies.

## Acknowledgements

This project was supported by the *Compte d’Affectation Spéciale et Développement Agricole et Rural* (CASDAR), the *Programme Complémentaire Intercéréales* and the ANSES Aflafrance project. We thank Olivier Puel and Sophie Lorber for providing several fungal strains. We also thank the laboratory team warmly for their advice and help with experiments. We also thank the team responsible for preparing the maize flour samples. We thank Marie Foulongne-Oriol for critical reading of the manuscript and valuable feedback.

## Data availability statement

The data that support the findings of this study are available on request from the corresponding author. The data are not publicly available due to privacy or ethical restrictions.

